# Discovery of a novel coronavirus associated with the recent pneumonia outbreak in humans and its potential bat origin

**DOI:** 10.1101/2020.01.22.914952

**Authors:** Peng Zhou, Xing-Lou Yang, Xian-Guang Wang, Ben Hu, Lei Zhang, Wei Zhang, Hao-Rui Si, Yan Zhu, Bei Li, Chao-Lin Huang, Hui-Dong Chen, Jing Chen, Yun Luo, Hua Guo, Ren-Di Jiang, Mei-Qin Liu, Ying Chen, Xu-Rui Shen, Xi Wang, Xiao-Shuang Zheng, Kai Zhao, Quan-Jiao Chen, Fei Deng, Lin-Lin Liu, Bing Yan, Fa-Xian Zhan, Yan-Yi Wang, Geng-Fu Xiao, Zheng-Li Shi

**Author notes:** These authors contributed equally (Peng Zhou, Xing-Lou Yang, Xian-Guang Wang).

## Abstract

Since the SARS outbreak 18 years ago, a large number of severe acute respiratory syndrome related coronaviruses (SARSr-CoV) have been discovered in their natural reservoir host, bats^1-4^. Previous studies indicated that some of those bat SARSr-CoVs have the potential to infect humans^5-7^. Here we report the identification and characterization of a novel coronavirus (nCoV-2019) which caused an epidemic of acute respiratory syndrome in humans, in Wuhan, China. The epidemic, started from December 12^th^, 2019, has caused 198 laboratory confirmed infections with three fatal cases by January 20^th^, 2020. Full-length genome sequences were obtained from five patients at the early stage of the outbreak. They are almost identical to each other and share 79.5% sequence identify to SARS-CoV. Furthermore, it was found that nCoV-2019 is 96% identical at the whole genome level to a bat coronavirus. The pairwise protein sequence analysis of seven conserved non-structural proteins show that this virus belongs to the species of SARSr-CoV. The nCoV-2019 virus was then isolated from the bronchoalveolar lavage fluid of a critically ill patient, which can be neutralized by sera from several patients. Importantly, we have confirmed that this novel CoV uses the same cell entry receptor, ACE2, as SARS-CoV.

---

Coronavirus has caused two large-scale pandemic in the last two decades, SARS and MERS (Middle East respiratory syndrome)^8,9^. It was generally believed that SARSr-CoV, mainly found in bats, might cause future disease outbreak^10,11^. Here we report on a series of unidentified pneumonia disease outbreaks in Wuhan, Hubei province, central China (Extended Data Figure 1). Started from a local fresh seafood market, the epidemic has resulted in 198 laboratory confirmed cases with three death according to authorities so far^12^. Typical clinical symptoms of these patients are fever, dry cough, dyspnea, headache, and pneumonia. Disease onset may result in progressive respiratory failure due to alveolar damage and even death. The disease was determined as viral induced pneumonia by clinicians according to clinical symptoms and other criteria including body temperature rising, lymphocytes and white blood cells decreasing (sometimes normal for the later), new pulmonary infiltrates on chest radiography, and no obvious improvement upon three days antibiotics treatment. It appears most of the early cases had contact history with the original seafood market, and no large scale of human-to-human transmission was observed so far.

**Fig. 1.**
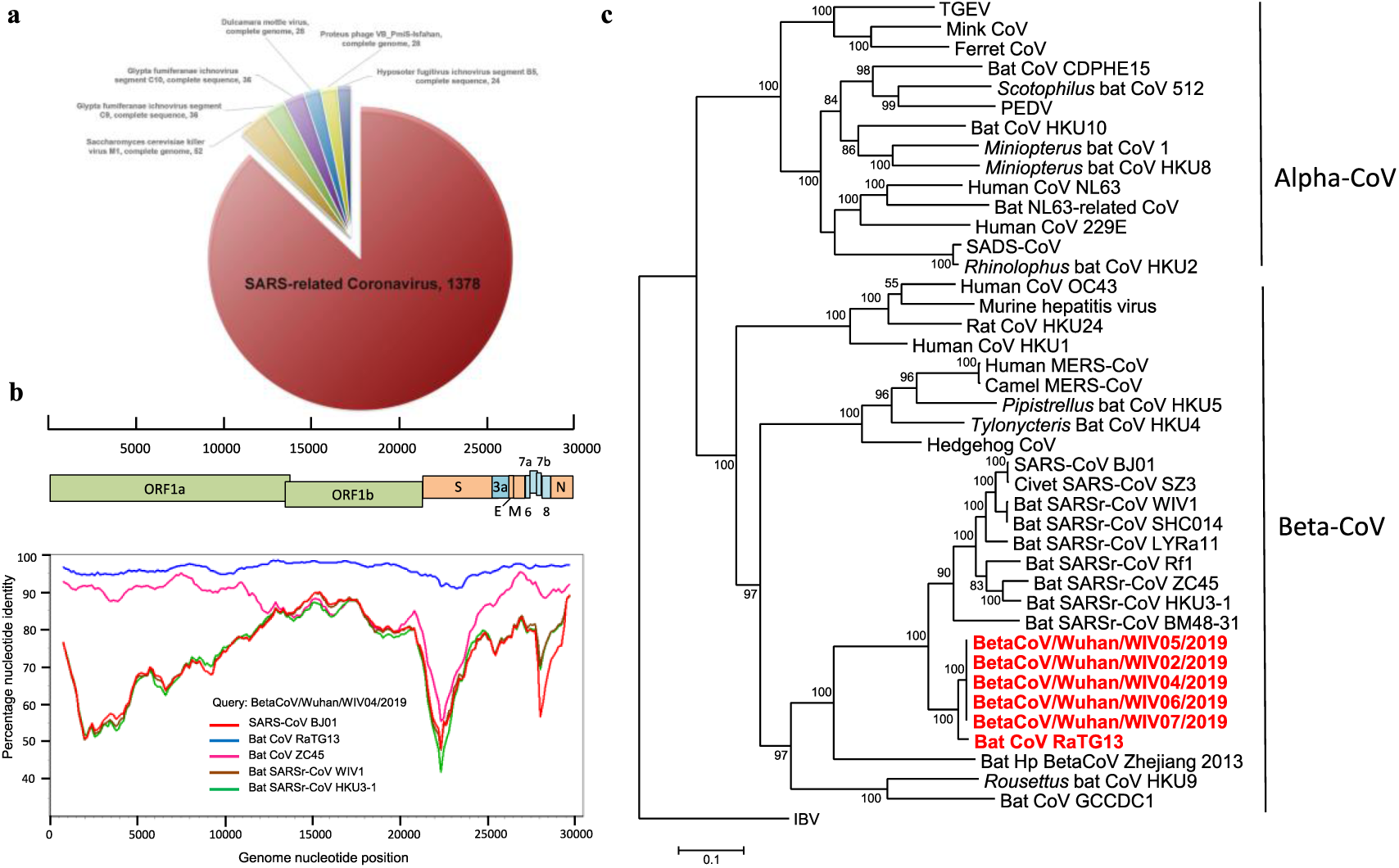
Genome characterization of nCoV-2019. **a**, pie chart showing metagenomics analysis of next-generation sequencing of bronchoalveolar lavage fluid from patient ICU06. **b**, Genomic organization of nCoV-2019 WIV04. **c**, Similarity plot based on the full-length genome sequence of nCoV-2019 WIV04. Full-length genome sequences of SARS-CoV BJ01, bat SARSr-CoV WIV1, bat coronavirus RaTG13 and ZC45 were used as reference sequences. **d**, Phylogenetic tree based on nucleotide sequences of complete ORF1b of coronaviruses. Software used and settings can be found in material and method section.

Samples from seven patients with severe pneumonia (six are seafood market peddlers or delivers), who were enrolled in intensive unit cares at the beginning of the outbreak, were sent to WIV laboratory for pathogen diagnosis (Extended Data Table 1). As a CoV lab, we first used pan-CoV PCR primers to test these samples^13^, considering the outbreak happened in winter and in a market, same environment as SARS. We found five PCR positive. A sample (WIV04) collected from bronchoalveolar lavage fluid (BALF) was analysed by metagenomics analysis using next-generation sequencing (NGS) to identify potential etiological agents. Of the 1582 total reads obtained after human genome filtering, 1378 (87.1%) matched sequences of SARSr-CoV (Fig. 1a). By *de novo* assembly and targeted PCR, we obtained a 29,891-bp CoV genome that shared 79.5% sequence identity to SARS-CoV BJ01 (GenBank accession number AY278488.2). This sequence has been submitted to GISAID (accession no. EPI_ISL_402124). Following the name by WHO, we tentatively call it novel coronavirus 2019 (nCoV-2019). Four more full-length genome sequences of nCoV-2019 (WIV02, WIV05, WIV06, and WIV07) (GISAID accession nos. EPI_ISL_402127-402130) that were above 99.9% identical to each other were subsequently obtained from other four patients (Extended Data Table 2).

The virus genome consists of six major open reading frames (ORFs) common to coronaviruses and a number of other accessory genes (Fig. 1b). Further analysis indicates that some of the nCoV-2019 genes shared less than 80% nt sequence identity to SARS-CoV. However, the seven conserved replicase domains in ORF1ab that were used for CoV species classification, are 94.6% aa sequence identical between nCoV-2019 and SARS-CoV, implying the two belong to same species (Extended Data Table 3).

We then found a short RdRp region from a bat coronavirus termed BatCoV RaTG13 which we previously detected in *Rhinolophus affinis* from Yunnan Province showed high sequence identity to nCoV-2019. We did full-length sequencing to this RNA sample. Simplot analysis showed that nCoV-2019 was highly similar throughout the genome to RaTG13 (Fig. 1c), with 96.2% overall genome sequence identity. The phylogenetic analysis also showed that RaTG13 is the closest relative of the nCoV-2019 and form a distinct lineage from other SARSr-CoVs (Fig. 1d). The receptor binding protein spike (S) gene was highly divergent to other CoVs (Extended Data Figure 2), with less than 75% nt sequence identity to all previously described SARSr-CoVs except a 93.1% nt identity to RaTG13 (Extended Data Table 3). The S genes of nCoV-2019 and RaTG13 S gene are longer than other SARSr-CoVs. The major differences in nCoV-2019 are the three short insertions in the N-terminal domain, and four out of five key residues changes in the receptor-binding motif, in comparison with SARS-CoV (Extended Data Figure 3). The close phylogenetic relationship to RaTG13 provides evidence for a bat origin of nCoV-2019.

**Fig. 2.**
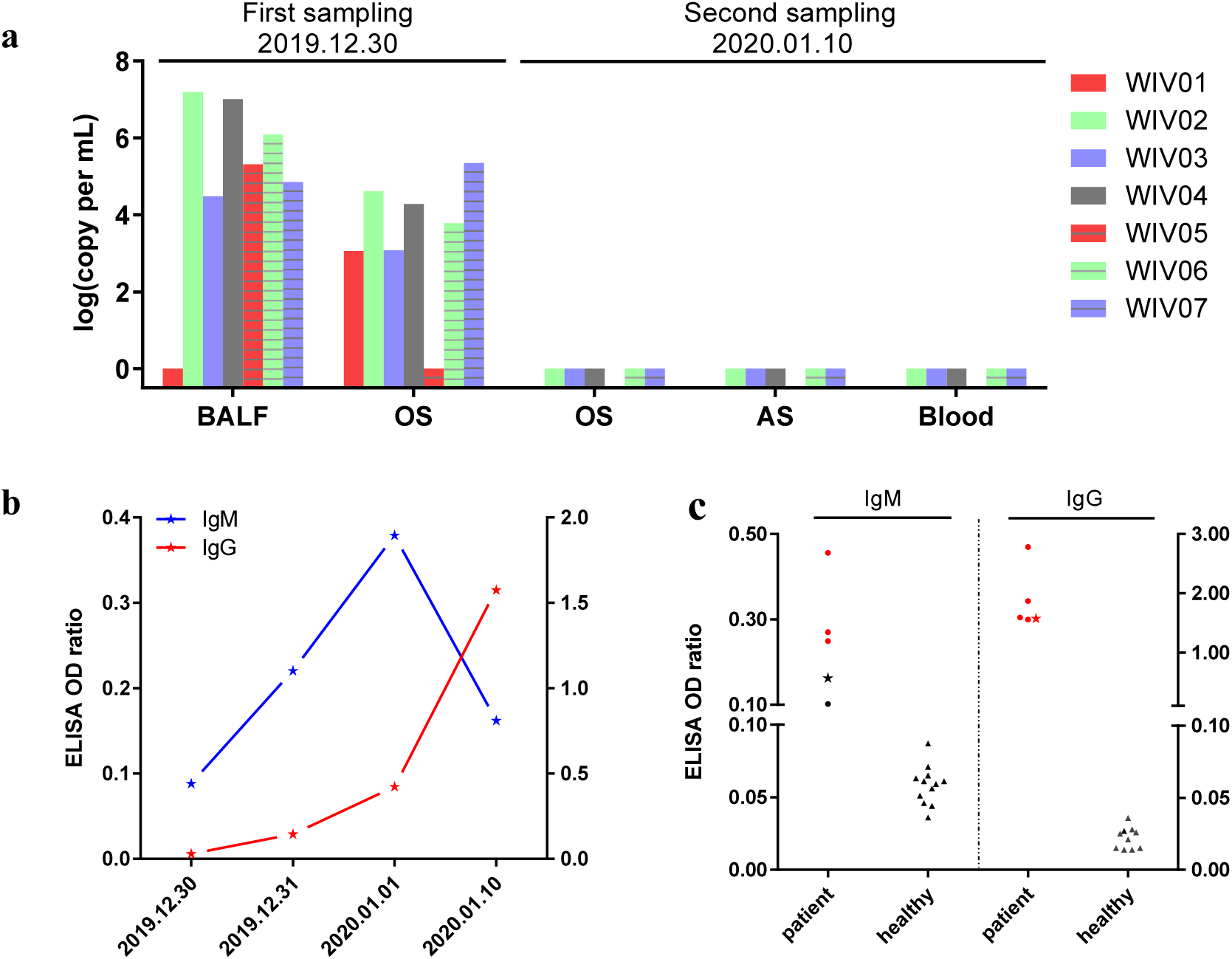
Molecular and serological investigation of patient samples. **a**, molecular detection of nCoV-2019 in seven patients during two times of sampling. Patient information can be found in Extended Data Table 1 and 2. Details on detection method can be found in material and methods. BALF, bronchoalveolar lavage fluid; OS, oral swab; AS, anal swab. **b**, dynamics of nCoV-2019 antibodies in one patient who showed sign of disease on 2019.12.23 (ICU-06). **c**, serological test of nCoV-2019 antibodies in five patients (more information can be found in Extended Data Table 2). Star indicates data collected from patient ICU-06 on 2020.01.10. For b and c, cut-off was set up as 0.2 for IgM test and 0.3 for IgG test, according to healthy controls.

**Fig. 3.**
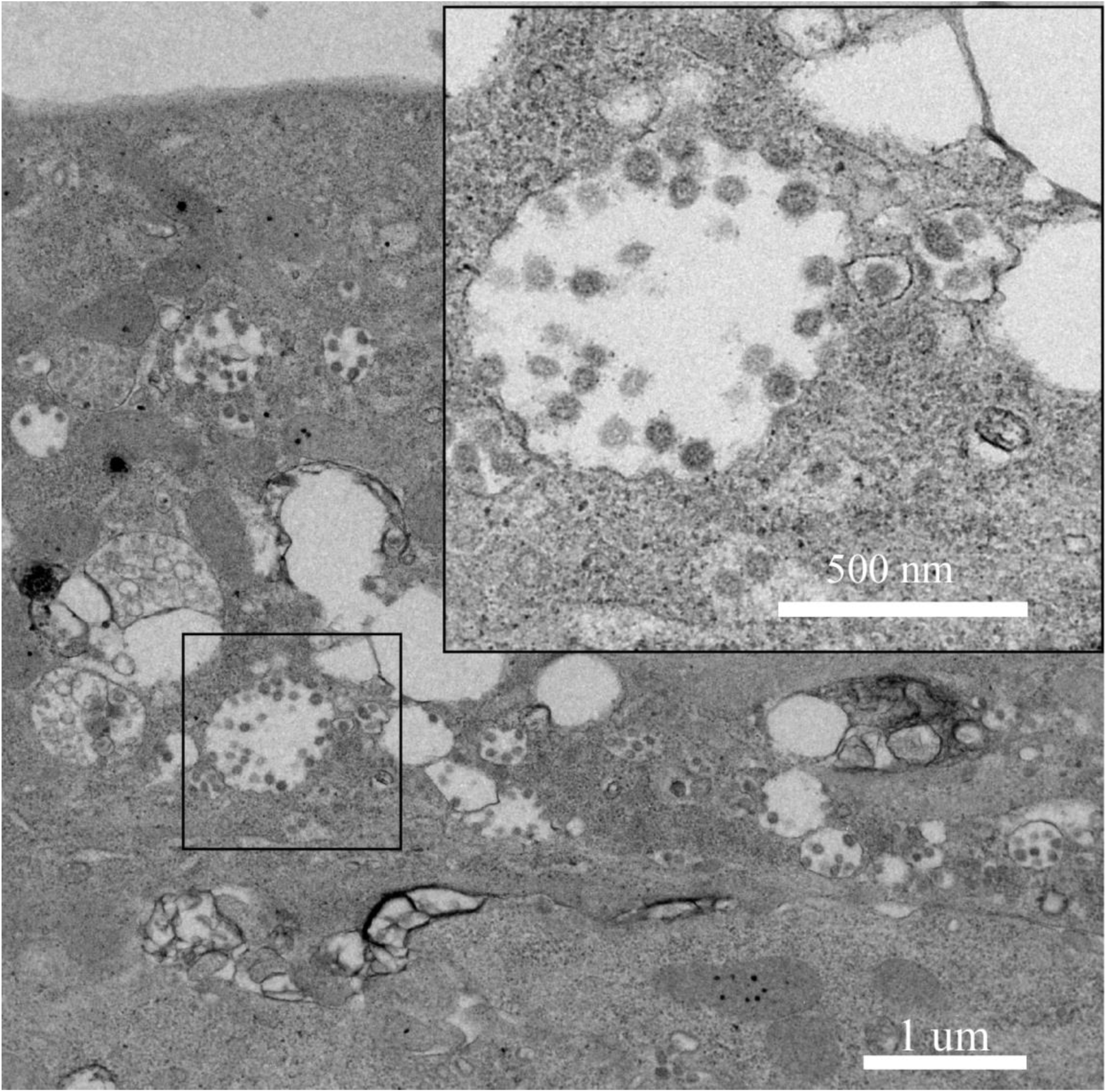
Virions. **a**, viral particles in the ultrathin sections under electron microscope at 200 kV, sample from viral infected Vero E6 cells

We rapidly developed a qPCR detection based on the receptor-binding domain of spike gene, the most variable region among genome (Fig. 1c). Our data show the primers could differentiate nCoV-2019 with all other human coronaviruses including bat SARSr-CoV WIV1, which is 95% identity to SARS-CoV (Extended Data Figure 4a and 4b). From the seven patients, we found nCoV-2019 positive in six BALF and five oral swab samples during the first sampling by qPCR and conventional PCR (Extended Data Figure 4c). However, we can no longer find viral positive in oral swabs, anal swabs, and blood from these patients during the second sampling (Fig. 2a). Based on these findings, we conclude that the disease should be transmitted through airway, yet we can’t rule out other possibilities if the investigation extended to include more patients.

**Fig. 4.**
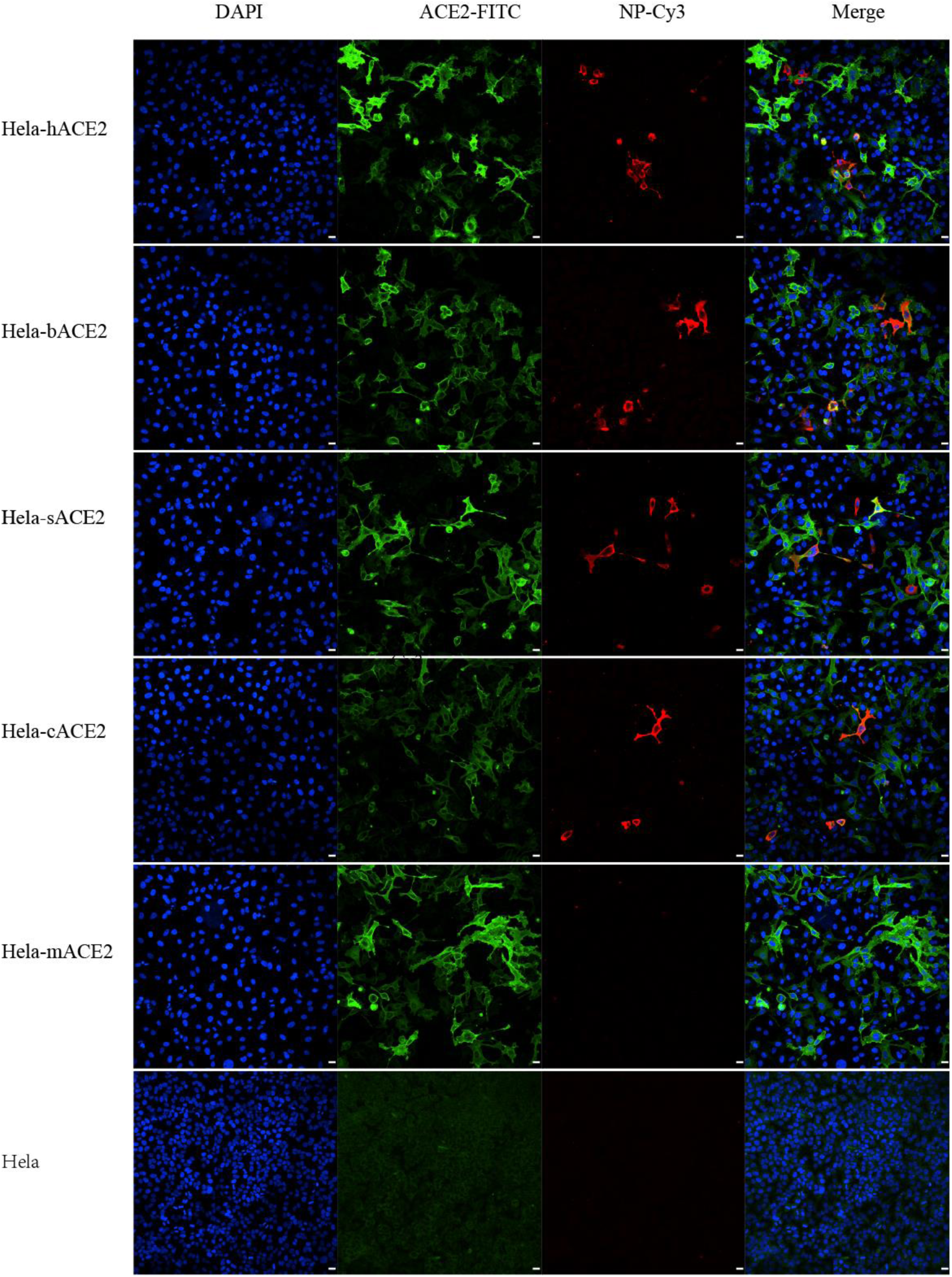
Analysis of nCoV-2019 receptor usage. Determination of virus infectivity in HeLa cells with or without the expression of ACE2. h, human; b, *Rhinolophus sinicus* bat; c, civet; s, swine (pig); m, mouse. ACE2 protein (green), viral protein (red) and nuclei (blue) was shown. Scale bar=10 um.

For serological detection of nCoV-2019, we used previously developed bat SARSr-CoV Rp3 nucleocapsid protein (NP) as antigen in IgG and IgM ELISA test, which showed no cross-reactivity against other human coronaviruses except SARSr-CoV^7^. As a research lab, we were only able to get five serum samples from the seven viral infected patients. We monitored viral antibody levels in one patient (ICU-06) at seven, eight, nine, and eighteen days after disease onset (Extended Data Table 2). A clear trend of IgG and IgM titre (decreased at the last day) increase was observed (Fig. 2b). For a second investigation, we tested viral antibody for five of the seven viral positive patients around twenty days after disease onset (Extended Data Table 1 and 2). All patient samples, but not samples from healthy people, showed strong viral IgG positive (Fig. 2b). We also found three IgM positive, indicating acute infection.

We then successfully isolated the virus (named nCoV-2019 BetaCoV/Wuhan/WIV04/2019), in Vero and Huh7 cells using BALF sample from ICU-06 patient. Clear cytopathogenic effects were observed in cells after three days incubation (Extended Data Figure 5a and 5b). The identity of the strain WIV04 was verified in Vero E6 cells by immunofluorescence microscopy using cross-reactive viral NP antibody (Extended Data Figure 5c and 5d), and by metagenomic sequencing, from which most of the reads mapped to nCoV-2019 (Extended Data Figure 5e and 5f). Viral partials in ultrathin sections of infected cells displayed typical coronavirus morphology under electron microscopy (Fig. 3). To further confirm the neutralization activity of the viral IgG positive samples, we conducted serum-neutralization assays in Vero E6 cells using the five IgG positive patient sera. We demonstrate that all samples were able to neutralize 120 TCID50 nCoV-2019 at a dilution of 1:40-1:80. We also show that this virus could be cross-neutralized by horse anti-SARS-CoV serum at dilutions 1:80, further confirming the relationship of the two viruses (Extended Data Table 4).

Angiotensin converting enzyme II (ACE2) was known as cell receptor for SARS-CoV^14^. To determine whether nCoV-2019 also use ACE2 as a cellular entry receptor, we conducted virus infectivity studies using HeLa cells expressing or not expressing ACE2 proteins from humans, Chinese horseshoe bats, civet, pig, and mouse. We show that nCoV-2019 is able to use all but mouse ACE2 as an entry receptor in the ACE2-expressing cells, but not cells without ACE2, indicating which is likely the cell receptor of nCoV-2019 (Fig. 4). We also proved that nCoV-2019 does not use other coronavirus receptors, aminopeptidase N and dipeptidyl peptidase 4 (Extended Data Figure 6).

The study provides the first detailed report on nCoV-2019, the likely etiology agent responsible for ongoing acute respiratory syndrome epidemic in Wuhan, central China. Viral specific nucleotide positive and viral protein seroconversion observed in all patients tested provides evidence of an association between the disease and the presence of this virus. However, there are still many urgent questions to be answered. We need more clinical data and samples to confirm if this virus is indeed the etiology agent for this epidemic. In addition, we still don’t know if this virus continue evolving and become more transmissible between human-to-human. Moreover, we don’t know the transmission routine of this virus among hosts yet. We showed viral positive in oral swabs, implying nCoV-2019 may be transmitted through airway. However, this needs to be confirmed by extending detection range. Finally, based on our results, it should be expected and worth to test if ACE2 targeting or SARS-CoV targeting drugs can be used for nCoV-2019 patients. At this stage, we know very little about the virus, including basic biology, animal source or any specific treatment. The almost identical sequences of this virus in different patients imply a probably recent introduction in humans, thus future surveillance on viral mutation and transmission ability and further global research attention are urgently needed.

## ACKNOWLEDGEMENTS

We thank the Pei Zhang and An-na Du from WIV core facility and technical support for their help with producing TEM micrographs. This work was jointly supported by the Strategic Priority Research Program of the Chinese Academy of Sciences (XDB29010101 to ZLS and XDB29010104 to PZ), China Natural Science Foundation for excellent scholars (81822028 to PZ, 31770175 to ZLS and 31800142 to BH), Mega-Project for Infectious Disease from Minister of Science and Technology of the People’s Republic of China (2020ZX09201001 to DYZ and 2018ZX10305409-004-001 to PZ), Youth innovation promotion association of CAS (2019328 to XLY).

## AUTHOR CONTRIBUTIONS

Z.L.S., P.Z., Y.Y.W., and G.F.X. conceived the study. G.S.W., C.L.H., H.D.C., F.D., Q.J.C., F.X.Z., and LLL., collected patient samples. X.L.Y., B.Y., W.Z., B.L., J.C., X.S.Z., Y.L., H.G., R.D.J., M.Q.L., Y. Chen, X.W., X.R.S., and K.Z. performed qPCR, serology, and virus culturing. L.Z., Y.Z., H.R.S., and B.H. performed genome sequencing and annotations. The authors declare no competing financial interests. Correspondence and requests for materials should be addressed to ZLS.

**Supplementary Information is available in the online version of the paper.**

## METHODS

### Sample collection

Human samples, including oral swabs, anal swabs, blood, and BALF samples were collected by Jinyintan hospital (Wuhan) with the consent from all patients. Patients were sampled without gender or age preference unless where indicated. For swabs, 1.5 ml DMEM+2% FBS medium was added each tube. Supernatant was collected after 2500 rpm, 60 s vortex and 15-30 min standing. Supernatant from swabs or BALF (no pretreatment) was added to either lysis buffer for RNA extraction or to viral transport medium (VTM) for virus isolation. VTM composed of Hank’s balanced salt solution at pH7.4 containing BSA (1%), amphotericin (15 μg/ml), penicillin G (100 units/ml), and streptomycin (50 μg/ml). Serum was separated by centrifugation at 3,000 *g* for 15 min within 24 h of collection, followed by 56 °C 30 min inactivation, and then stored at 4 °C until use.

### Virus isolation, cell infection, electron microscope and neutralization assay

The following cells were used for virus isolation in this study: Vero, Vero E6, and Huh7 that were cultured in DMEM +10% FBS. A list of cells were used for susceptibility test (Extended Data Fig. 6). All cell lines were tested free of mycoplasma contamination, applied to species identification and authenticated by microscopic morphologic evaluation. None of cell lines was on the list of commonly misidentified cell lines (by ICLAC).

Cultured cell monolayers were maintained in their respective medium. PCR-positive BALF sample from ICU-06 patient was spin at 8,000 g for 15 min, filtered and diluted 1:2 with DMEM supplied with 16 μg/ml trypsin before adding to cells. After incubation at 37 °C for 1 h, the inoculum was removed and replaced with fresh culture medium containing antibiotics (below) and 16 μg/ml trypsin. The cells were incubated at 37 °C and observed daily for cytopathic effect (CPE). The culture supernatant was examined for presence of virus by qRT-PCR developed in this study, and cells were examined by immunofluorescent using SARSr-CoV Rp3 NP antibody made in house (1:100). Penicillin (100 units/ml) and streptomycin (15 μg/ml) were included in all tissue culture media.

The Vero E6 cells were infected with new virus at MOI of 0.5 and harvested 48 hpi. Cells were fixed with 2.5% (wt/vol) glutaraldehyde and 1% osmium tetroxide, and then dehydrated through a graded series of ethanol concentrations (from 30 to 100%), and embedded with epoxy resin. Ultrathin sections (80 nm) of embedded cells were prepared, deposited onto Formvar-coated copper grids (200 mesh), stained with uranyl acetate and lead citrate, then observed under 200 kV Tecnai G2 electron microscope.

The virus neutralization test was carried out in a 48-well plate. The patient serum samples were heat-inactivated by incubation at 56 °C for 30 min before use. The serum samples (5 µL) were diluted to 1:10, 1:20, 1:40 or 1:80, and then an equal volume of virus stock was added and incubated at 37 °C for 60 min in a 5% CO2 incubator. Diluted horse anti SARS-CoV serum or serum samples from healthy people were used as control. After incubation, 100 µL mixtures were inoculated onto monolayer Vero E6 cells in a 48-well plate for 1 hour. Each serum were repeated triplicate. After removing the supernatant, the plate was washed twice with DMEM medium. Cells were incubated with DMEM supplemented with 2% FBS for 24 hours. Then the cells were fixed with 4% formaldehyde. And the virus were detected using SL-CoV Rp3 NP antibody followed by Cy3-conjugated mouse anti-rabbit IgG. Nuclei were stained with DAPI. Infected cell number was counted by high-content cytometers.

### RNA extraction and PCR

Whenever commercial kits were used, manufacturer’s instructions were followed without modification. RNA was extracted from 200 μl of samples with the High Pure Viral RNA Kit (Roche). RNA was eluted in 50 μl of elution buffer and used as the template for RT-PCR.

For qPCR analysis, primers based on nCoV-2019 S gene was designed: RBD-qF1: 5’-CAATGGTTTAACAGGCACAGG-3’; RBD-qR1: 5’-CTCAAGTGTCTGTGGATCACG-3’. RNA extracted from above used in qPCR by HiScript^®^ II One Step qRT-PCR SYBR^®^ Green Kit (Vazyme Biotech Co.,Ltd). Conventional PCR test was also performed using the following primer pairs: ND-CoVs-951F TGTKAGRTTYCCTAAYATTAC; ND-CoVs-1805R ACATCYTGATANARAACAGC^13^. The 20 μl qPCR reaction mix contained 10 μl 2× One Step SYBR Green Mix, 1 μl One Step SYBR Green Enzyme Mix, 0.4 μl 50 × ROX Reference Dye 1, 0.4 μl of each primer (10 uM) and 2 μl template RNA. Amplification was performed as follows: 50 °C for 3 min, 95 °C for 30 s followed by 40 cycles consisting of 95 °C for 10 s, 60 °C for 30 s, and a default melting curve step in an ABI 7700 machine.

### Serological test

In-house anti-SARSr-CoV IgG and IgM ELISA kits were developed using SARSr-CoV Rp3 NP as antigen, which shared above 90% amino acid identity to all SARSr-CoVs^2^. For IgG test, MaxiSorp Nunc-immuno 96 well ELISA plates were coated (100 ng/well) overnight with recombinant NP. Human sera were used at 1:20 dilution for 1 h at 37 °C. An anti-Human IgG-HRP conjugated monoclonal antibody (Kyab Biotech Co., Ltd, Wuhan, China) was used at a dilution of 1:40000. The OD value (450–630) was calculated. For IgM test, MaxiSorp Nunc-immuno 96 wellELISA plates were coated (500 ng/well) overnight with anti-human IgM (µ chain). Human sera were used at 1:100 dilution for 40 min at 37 °C, followed by anti-Rp3 NP-HRP conjugated (Kyab Biotech Co., Ltd, Wuhan, China) at a dilution of 1:4000. The OD value (450–630) was calculated.

### Examination of ACE2 receptor for nCoV-2019 infection

HeLa cells transiently expressing ACE2 were prepared by a lipofectamine 3000 system (Thermo Fisher Scientific) in 96-well plate, with mock-transfected cells as controls. nCoV-2019 grown from Vero E6 cells was used for infection at multiplicity of infection 0.05. Same for testing of APN and DPP4. The inoculum was removed after 1 h absorption and washed twice with PBS and supplemented with medium. At 24 hpi, cells were washed with PBS and fixed with 4% formaldehyde in PBS (pH 7.4) for 20 min at room temperature. ACE2 expression was detected using mouse anti-S tag monoclonal antibody followed by FITC-labelled goat anti-mouse IgG H&L (Abcam, ab96879). Viral replication was detected using rabbit antibody against the Rp3 NP protein (made in house, 1:100) followed by cyanin 3-conjugated goat anti-rabbit IgG (1:50, Abcam, ab6939). Nucleus was stained with DAPI (Beyotime). Staining patterns were examined using the FV1200 confocal microscopy (Olympus).

### High throughput sequencing, pathogen screening and genome assembly

Samples from patient BALF or from virus culture supernatant were used for RNA extraction and next-generation sequencing using Illumina MiSeq 3000 sequencer. Metagenomic analysis was carried out mainly base on the bioinformatics platform MGmapper (PE_2.24 and SE_2.24). The raw NGS reads were firstly processed by Cutadapt (v1.18) with minimum read length of 30bp. BWA (v0.7.12-r1039) was utilized to align reads to local database with a filter hits parameter at 0.8 FMM value and minimum alignment score at 30. Parameters for post-processing of assigned reads was set with minimum size normalized abundance at 0.01, minimum read count at 20 and other default parameters. A local nucleic acid database for human and mammals was employed to filter reads of host genomes before mapping reads to virus database. The results of metagenomic analysis were displayed through pie charts using WPS Office 2010. NGS reads were assembled into genomes using Geneious (v11.0.3) and MEGAHIT (v1.2.9). PCR and Sanger sequencing was performed to fill gaps in the genome. 5’-RACE was performed to determine the 5’-end of the genomes using SMARTer RACE 5’/3’ Kit (Takara). Genomes were annotated using Clone Manager Professional Suite 8 (Sci-Ed Software).

### Phylogenetic analysis

Routine sequence management and analysis was carried out using DNAStar. Sequence alignment and editing were conducted using ClustalW and GeneDoc. Maximum Likelihood phylogenetic trees based on nucleotide sequences of full-length ORF1b and S genes were constructed using the Jukes-Cantor model with bootstrap values determined by 1000 replicates in the MEGA6 software package.

## Data Availability statement

Sequence data that support the findings of this study have been deposited in GISAID with the accession no. EPI_ISL_402124 and EPI_ISL_402127-402130.

