## Extended Data Figures 1-6 and Tables 1-4 for "Discovery of a novel coronavirus associated with the recent pneumonia outbreak in humans and its potential bat origin"

*These authors contributed equally.

**Extended Data Fig. 1 |** Map of Wuhan. Wuhan, located in central China Hubei province (circled), has more than 11 million citizens.

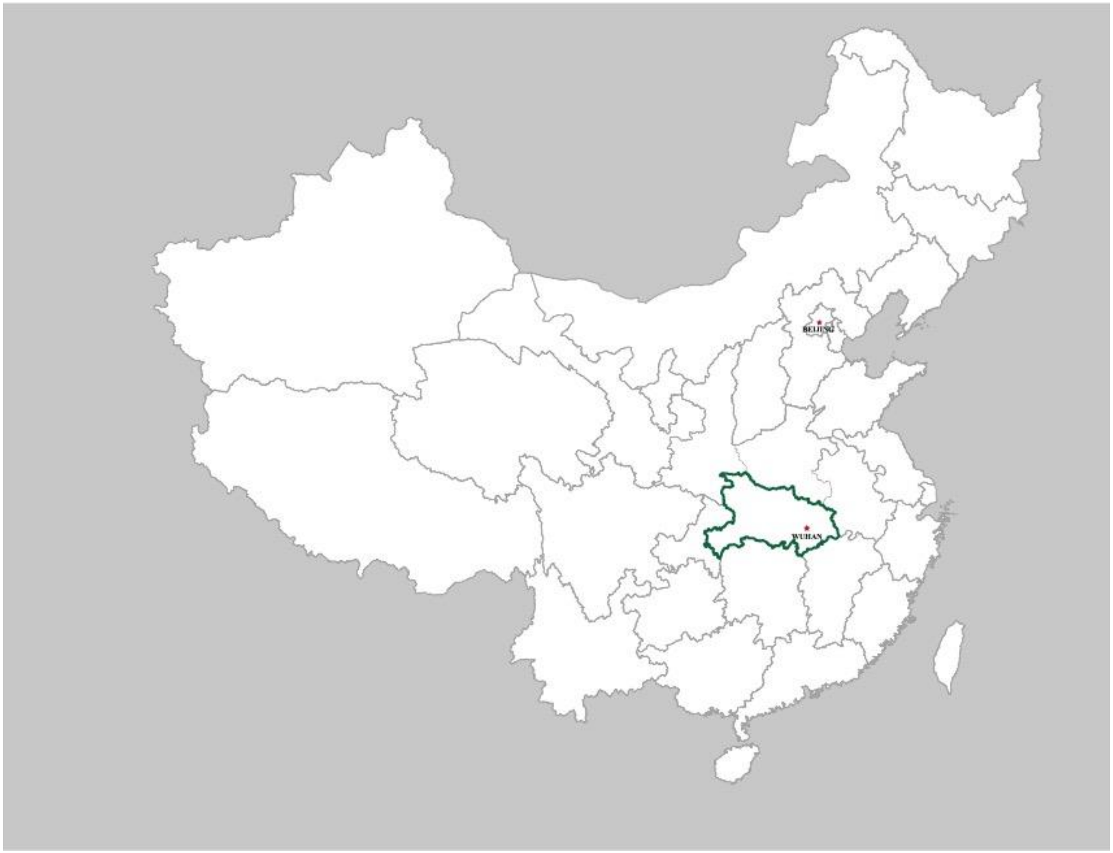

**Extended Data Fig. 2 |** Phylogenetic tree base on the complete S gene sequence. nCoV-2019 and bat CoV RaTG13 are in bold. R.s, *Rhinolophus sinicus*; R.a, *Rhinolophus affinis*; R.f, *Rhinolophus ferrumequinum*; R.m, *Rhinolophus macrotis*; R.b, *Rhinolophus blasii*. Bat CoV HKU9-1 was used as outgroup. The trees were constructed by the maximum likelihood method using the Jukes-Cantor model with bootstrap values determined by 1000 replicates. Bootstraps > 50% are shown.

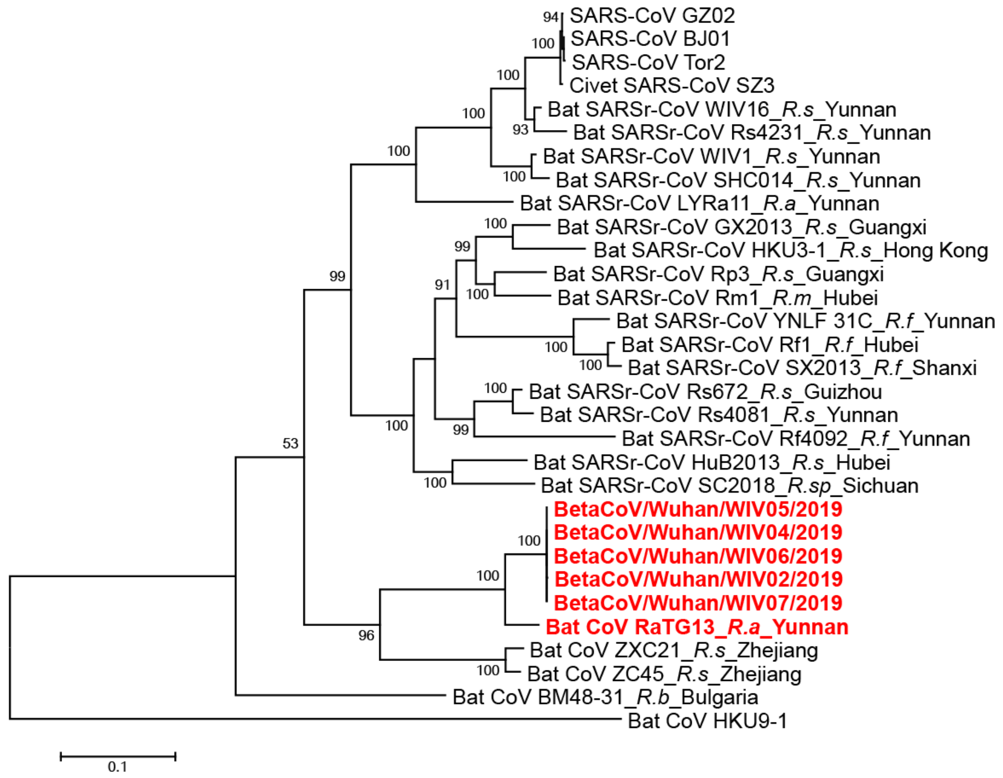

**Extended Data Fig. 3 |** Amino acid sequence alignment of the S1 protein of the nCoV-2019 with SARS-CoV and selected bat SARSr-CoVs. The receptor-binding motif of SARS-CoV and the homologous region of other coronaviruses are indicated by the red box. The key amino acid residues involved in the interaction with human ACE2 are numbered on top of the aligned sequences. The short insertions in the N-terminal domain of the novel coronavirus are indicated by the blue boxes. Bat CoV RaTG13 was identified from *R. affinis* in Yunnan Province. Bat CoV ZC45 was identified from *R. sinicus* in Zhejiang Province.

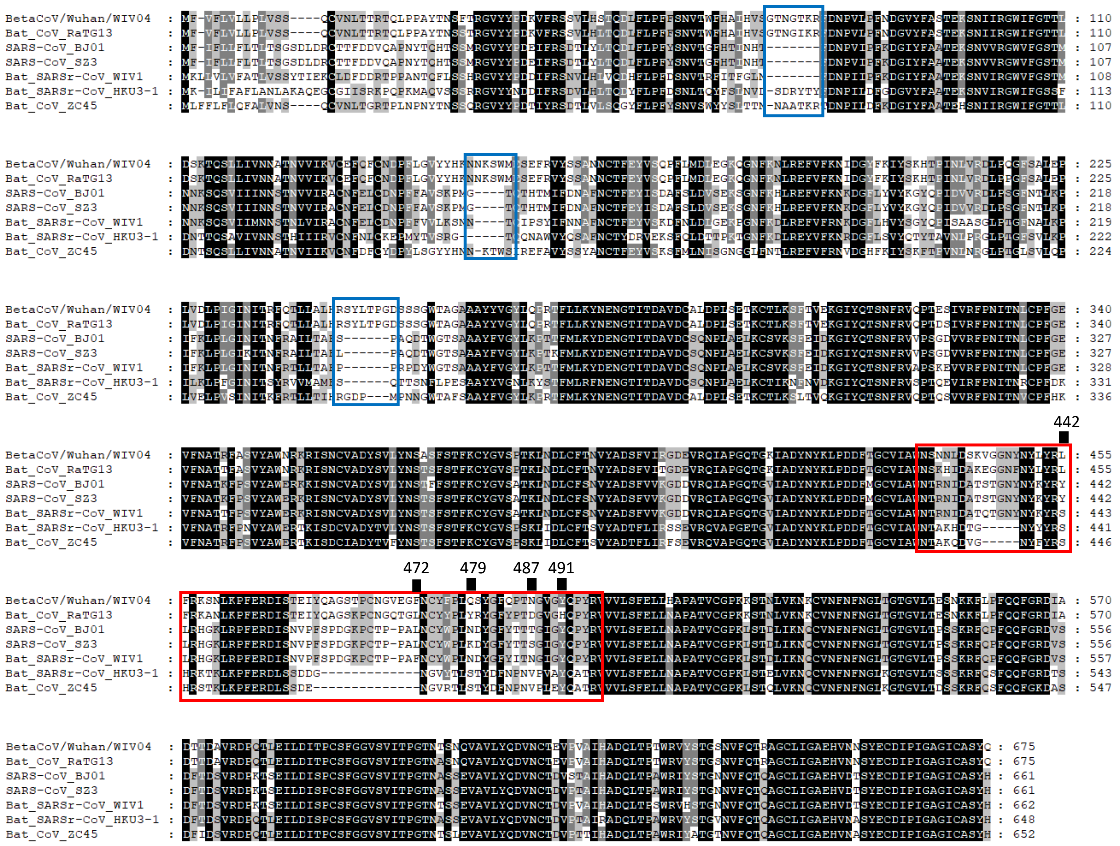

**Extended Data Fig. 4 |** Molecular detection method set up for nCoV-2019. **a**, molecular detection using conventional PCR. Primer sequence can be found in material and methods. **b**, standard curve for qPCR primers. PCR product of spike gene that was serial diluted to $10^8$ to $10^1$ (from left to right) was used as template. Primer sequence and experiment condition can be found in material and methods. **c**, specificity of qPCR primers. Nucleotide samples from the indicated pathogens were used.

**a**

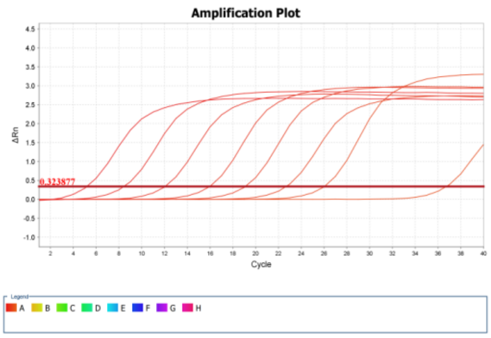


**c**

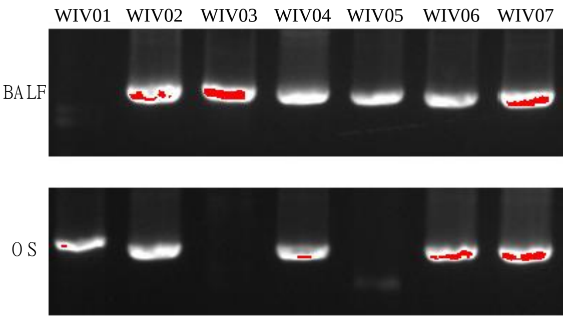


**b**

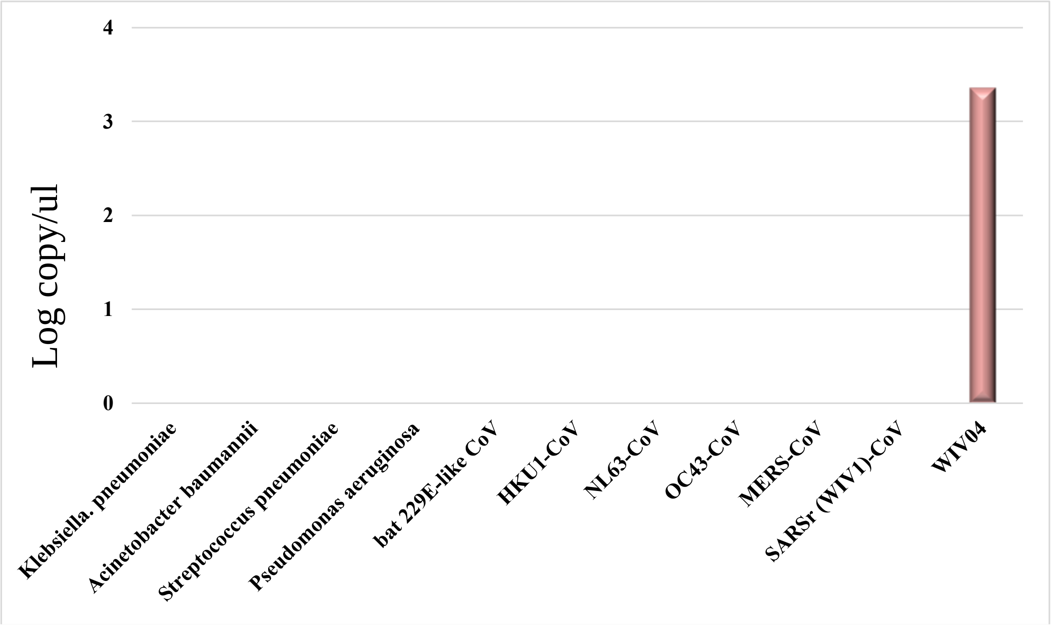

**Extended Data Fig. 5 |** Isolation and antigenic characterization of nCoV-2019. Vero E6 cells are shown at 24 hours post infection with mock (**a**) or nCoV-2019 (**b**). **c** and **d** are mock or nCoV-2019 infected samples stained with rabbit serum raised against recombinant SARSr-CoV Rp3 N protein (red) and DAPI (blue). The experiment was conducted two times independently with similar results. **e** and **f,** pie charts illustrating ratio of reads number related to nCoV-2019 among total viral related reads in metagenomics analysis of Vero (**e**) and Huh7 (**f**) cell culture supernatant.

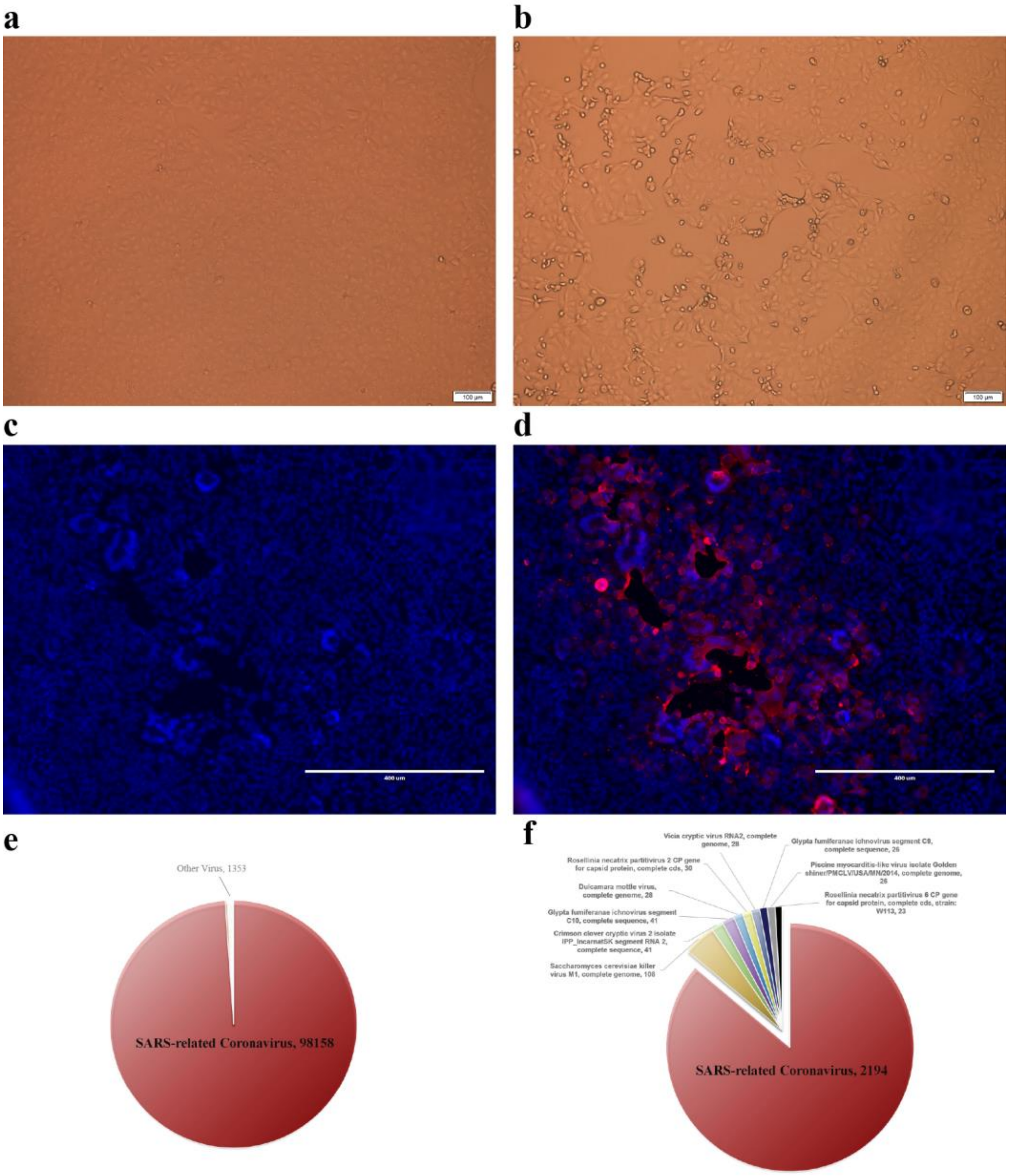

**Extended Data Fig. 6 | Analysis of nCoV-2019 receptor usage.** Determination of virus infectivity in HeLa cells with or without the expression of human APN and DPP4. ACE2 protein (green), viral protein (red) and nuclei (blue) were shown. Scale bar=10 um.

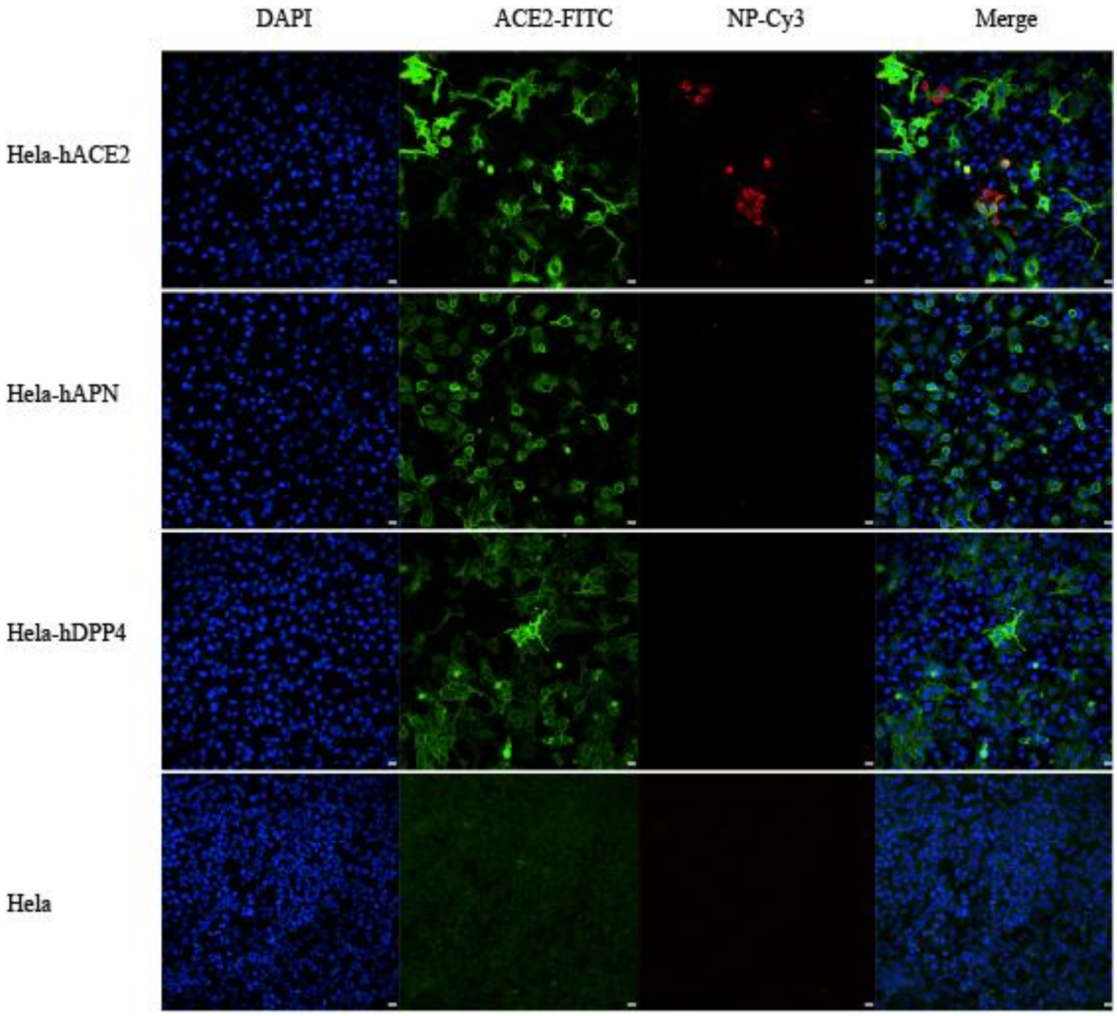

**Extended Data Table 1 |** Patient information and their diagnosis history (some records are missing). All patients are fresh seafood market peddlers or deliverymen except ICU-01, whose contact history is unclear. All patients were in intensive care unit (ICU) during the first investigation, and now in stable condition. Blood IgM tests have been performed for the following respiratory pathogens for all patients: legionella pneumophilia, mycoplasma pneumoniae, chlamydia pneumoniae, respiratory syncytial virus, adenovirus, rickettsia, influenza A virus, influenza B virus, parainfluenza virus.

| Patient No. | Gender | Age | Date of Onset | Date of Admission | Symptoms When Admitted | Current Status (2020.01.13) | Diagnosis history |
|---|---|---|---|---|---|---|---|
| ICU-01* | Male | 62 | 2019.12.12 | 2019.12.27 | fever | recover, discharged | negative |
| ICU-04 | Male | 32 | 2019.12.19 | 2019.12.29 | fever, cough, dyspnea | fever, intermittent cough | negative |
| ICU-05 | Male | 40 | 2019.12.17 | 2019.12.27 | fever (38 $^{o}$C), expectoration, malaise, dyspnea | fever, malaise, intermittent cough | AdV (IgM) |
| ICU-06 | Female | 49 | 2019.12.23 | 2019.12.27 | fever (37.9 $^{o}$C), palpitation | fever, malaise, cough | Coronavirus (nt) |
| ICU-08 | Female | 52 | 2019.12.22 | 2019.12.29 | fever (38.5 $^{o}$C), expectoration, malaise, dyspnea | recover, discharged | Streptococcus pneumoniae (nt) |
| ICU-09 | Male | 40 | 2019.12.22 | 2019.12.28 | fever (38.5 $^{o}$C), expectoration | fever (38.5 $^{o}$C), malaise, expectoration, dizziness | negative |
| ICU-10 | Male | 56 | 2019.12.20 | 2019.12.20 | fever, dyspnea, chest tightness | fever, malaise, cough, dyspnea | negative |

**Extended Data Table 2 |** Laboratory detection results. Samples from two patients (ICU-01 and ICU-08) were not available during the second investigation. They have been discharged from hospital. We did serial test for ICU-06 patient at the following date: 19.12.30, 19.12.31, 20.01.01 and 20.01.10, corresponding to seven, eight, nine and eighteen days upon disease onset (19.12.23). Table shows molecular and serological (IgM and IgG) detection results for nCoV-2019.

| | | First sampling-2019.12.30 | | | Second sampling-2020.01.10 | | | |
|---|---|---|---|---|---|---|---|---|
| Patient No. | Test No. | BALF | Oral Swab | Blood (Ab) | Oral Swab | Anal Swab | Blood (PCR) | Blood (Ab) |
| ICU-01 | WIV01 | - | + | NA | NA | NA | NA | NA |
| ICU-04 | WIV02# | + | + | NA | - | - | - | + |
| ICU-05 | WIV03 | + | + | NA | - | - | - | + |
| ICU-06 | WIV04#* | + | + | + | - | - | - | + |
| ICU-08 | WIV05# | + | - | NA | NA | NA | NA | NA |
| ICU-09 | WIV06# | + | + | NA | - | - | - | + |
| ICU-10 | WIV07# | + | + | NA | - | - | - | + |

**Extended Data Table 3 |** Genomic comparison of nCoV-2019 WIV04 with SARS-CoVs and bat SARSr-CoVs.

| | Sequence identities with SARS-CoVs & bat SARSr-CoVs (nt/aa %) | | | | | | | | | | | |
|---|---|---|---|---|---|---|---|---|---|---|---|---|
| | Full-length genome | ORF1a | ORF1b | S | ORF3a | E | M | ORF6 | ORF7a | ORF7b | ORF8 | N |
| SARS-CoV GZ02 | 79.6 | 76.0/80.9 | 86.2/95.7 | 73.4/77.0 | 75.6/73.4 | 94.7/96.0 | 85.4/90.5 | 76.3/68.9 | 82.8/86.0 | 84.8/81.4 | 52.0/31.6 | 87.7/91.2 |
| SARS-CoV BJ01 | 79.6 | 76.0/80.8 | 86.2/95.7 | 73.4/76.9 | 75.3/72.6 | 94.7/96.0 | 85.6/90.5 | 75.8/67.2 | 82.8/86.0 | 84.8/81.4 | 51.1/- | 88.8/91.2 |
| SARS-CoV Tor2 | 79.6 | 76.0/80.9 | 86.2/95.8 | 73.4/76.7 | 75.4/72.6 | 94.7/96.0 | 85.6/90.5 | 76.3/68.9 | 82.8/86.0 | 84.8/81.4 | 51.1/- | 88.8/91.2 |
| SARS-CoV SZ3 | 79.6 | 76.0/81.0 | 86.2/95.8 | 73.4/76.9 | 75.4/72.6 | 94.7/96.0 | 85.3/90.0 | 76.3/68.9 | 82.8/86.0 | 84.8/81.4 | 52.3/31.6 | 88.8/91.2 |
| SARS-CoV PC4-227 | 79.5 | 76.0/80.8 | 86.1/95.6 | 73.4/76.7 | 75.5/72.6 | 94.7/96.0 | 85.1/90.0 | 75.8/68.9 | 82.8/86.0 | 84.8/81.4 | 52.3/- | 88.5/90.7 |
| Bat SARr-CoV RaTG13 | 96.2 | 96.0/98.0 | 97.3/99.3 | 93.1/97.7 | 96.3/97.8 | 99.6/100 | 95.5/99.6 | 98.4/100 | 95.6/97.5 | 99.2/97.7 | 97.0/95.0 | 96.9/99.0 |
| Bat SARr-CoV WIV1 | 79.7 | 76.0/80.7 | 85.9/95.8 | 73.4/77.6 | 76.1/74.5 | 95.6/96.0 | 84.8/90.0 | 78.0/73.8 | 85.0/88.4 | 85.6/83.7 | 65.8/57.9 | 88.5/90.9 |
| Bat SARSr-CoV WIV16 | 79.7 | 75.9/81.0 | 86.1/95.6 | 73.1/77.8 | 76.1/74.5 | 95.6/96.0 | 84.8/90.0 | 77.4/72.1 | 85.0/88.4 | 85.6/83.7 | 65.3/57.9 | 88.6/90.9 |
| Bat SARSr-CoV SHC014 | 79.6 | 75.9/80.9 | 85.9/95.8 | 73.3/77.7 | 76.1/74.5 | 95.6/96.0 | 84.8/90.0 | 78.0/70.5 | 84.4/88.4 | 85.6/83.7 | 65.8/58.7 | 88.6/90.9 |
| Bat SARSr-CoV Rs4231 | 79.7 | 76.0/81.0 | 86.2/95.8 | 72.9/77.5 | 75.8/74.1 | 94.3/94.7 | 84.4/90.0 | 76.9/67.2 | 85.0/88.4 | 85.6/83.7 | 65.3/57.9 | 88.8/91.4 |
| Bat SARSr-CoV YNLF31C | 79.0 | 75.7/80.6 | 85.8/95.7 | 71.4/75.5 | 75.0/71.2 | 94.3/96.0 | 84.7/89.6 | 76.9/70.5 | 83.1/87.6 | 86.4/83.7 | 50.3/31.3 | 88.3/90.5 |
| Bat SARSr-CoV LYRa11 | 79.6 | 75.8/80.6 | 85.7/95.6 | 73.9/77.3 | 77.2/76.3 | 94.7/94.7 | 85.1/90.0 | 78.5/70.5 | 82.0/85.1 | 81.1/81.4 | 66.7/57.9 | 89.0/91.6 |
| Bat SARSr-CoV ZC45 | 88.1 | 91.0/95.7 | 86.1/96.0 | 77.8/82.3 | 87.8/90.9 | 98.7/100 | 93.4/98.6 | 95.2/93.4 | 88.8/87.6 | 94.7/93.0 | 88.5/94.2 | 91.1/94.3 |
| Bat SARSr-CoV ZXC21 | 88.0 | 90.9/95.7 | 86.2/95.8 | 77.1/81.7 | 88.9/92.0 | 98.7/100 | 93.4/98.6 | 95.2/93.4 | 89.1/88.4 | 95.5/93.0 | 88.5/94.2 | 91.2/94.3 |
| Bat SARSr-CoV HuB2013 | 79.6 | 76.3/81.2 | 85.3/95.7 | 73.1/76.8 | 75.4/75.5 | 95.2/94.7 | 85.3/91.0 | 76.3/68.9 | 84.2/87.6 | 85.6/83.7 | 62.0/49.6 | 88.9/91.6 |
| Bat SARSr-CoV GX2013 | 79.1 | 75.9/80.8 | 86.0/95.9 | 73.1/77.1 | 75.6/73.0 | 94.7/96.0 | 84.8/91.4 | 77.4/68.9 | 85.0/86.8 | 84.1/79.1 | 51.4/31.6 | 87.9/90.2 |
| Bat SARSr-CoV SX2013 | 78.9 | 76.2/80.6 | 85.1/95.5 | 71.2/75.5 | 74.7/71.2 | 94.3/93.3 | 83.0/89.6 | 77.4/68.9 | 84.2/86.8 | 85.6/83.7 | 49.7/30.4 | 86.9/90.2 |
| Bat SARSr-CoV SC2018 | 79.4 | 75.8/80.7 | 85.5/95.2 | 72.7/76.4 | 75.0/71.2 | 94.3/96.0 | 84.7/90.0 | 80.0/71.8 | 85.2/87.6 | 84.8/83.7 | 66.1/55.4 | 88.2/91.2 |
| Bat SARSr-CoV Rs672 | 79.6 | 76.0/80.9 | 85.9/95.8 | 72.8/76.2 | 75.2/71.9 | 95.2/96.0 | 84.8/89.6 | 78.5/70.5 | 84.7/88.4 | 85.6/83.7 | 65.8/58.7 | 87.9/91.2 |
| Bat SARSr-CoV Rp3 | 79.5 | 75.9/80.5 | 86.0/95.7 | 73.1/77.2 | 74.9/74.8 | 95.2/96.0 | 85.1/90.0 | 76.9/68.9 | 83.9/89.3 | 84.8/83.7 | 66.4/56.2 | 88.4/90.7 |
| Bat SARSr-CoV Rf1 | 78.8 | 76.2/80.6 | 84.8/95.3 | 71.1/75.7 | 74.3/69.0 | 94.3/94.7 | 83.3/89.6 | 79.0/68.9 | 84.2/86.8 | 84.1/83.7 | 50.6/31.3 | 86.8/89.5 |
| Bat SARSr-CoV HKU3-1 | 79.4 | 76.1/80.9 | 84.9/95.1 | 73.4/77.9 | 75.8/73.4 | 95.2/96.0 | 84.7/91.0 | 75.3/67.2 | 85.0/89.3 | 84.1/79.1 | 66.4/57.0 | 88.3/90.0 |

**Extended Data Table 4 |** Virus neutralization test (VNT) of serum samples. Each serum sample was tested in triplicate. Two healthy people from Wuhan, five patient serum samples and a horse anti-SARS-CoV anti-serum were used. 120 $TCID_{50}$ virus was used each well. Serum samples were used in a dilution from 1:10, 1:20, 1:40 to 1:80.

| Samples | VNT titre for nCoV-2019 |
|---|---|
| Healthy people #1 from Wuhan | neg |
| Healthy people #2 from Wuhan | neg |
| Horse anti-SARS-CoV serum | >1:80 |
| WIV02 | >1:80 |
| WIV03 | 1:40 |
| WIV04 | >1:80 |
| WIV06 | >1:80 |
| WIV07 | >1:80 |
